# Electro-optical mechanically flexible coaxial microprobes for minimally invasive interfacing with intrinsic neural circuits

**DOI:** 10.1101/2020.09.16.300152

**Authors:** Spencer Ward, Conor Riley, Erin M. Carey, Jenny Nguyen, Sadik Esener, Axel Nimmerjahn, Donald J. Sirbuly

## Abstract

Central to advancing our understanding of neural circuits is the development of minimally invasive, multi-modal interfaces capable of simultaneously recording and modulating neural activity. Recent devices have focused on matching the mechanical compliance of tissue to reduce inflammatory responses^1,2^. However, reductions in the size of multi-modal interfaces are needed to further improve biocompatibility and long-term recording capabilities^1^. Here we demonstrate a multi-modal coaxial microprobe design with a minimally invasive footprint (8-12 μm diameter over millimeter lengths) that enables efficient electrical and optical interrogation of neural networks. In the brain, the probes allowed robust electrical measurement and optogenetic stimulation. Scalable fabrication strategies can be used with various electrical and optical materials, making the probes highly customizable to experimental requirements, including length, diameter, and mechanical properties. Given their negligible inflammatory response, these probes promise to enable a new generation of readily tunable multi-modal devices for minimally invasive interfacing with neural circuits.

---

Microelectrode recordings are the gold standard for measuring individual neurons’ activity at high temporal resolution in any nervous system region and central to defining the role of neural circuits in controlling behavior. Microelectrode arrays, such as the Utah^3^ or Michigan^4^ arrays, have allowed tracking of distributed neural activity with millisecond precision. However, their large footprint and rigidity lead to tissue damage and inflammation that hamper long-term recordings^1,5^. State of the art Neuropixel^6^ and carbon fiber probes^7^ have improved on these previous devices by increasing electrode density and reducing probe dimensions and rigidity. Although these probes have advanced the field of recordings, next-generation devices should enable targeted stimulation in addition to colocalized electrical recordings^1,2^. Optogenetic techniques enable high-speed modulation of cellular activity through targeted expression and activation of light-sensitive opsins^8^. However, given the strong light scattering and high absorption properties of neural tissue^9^ optogenetic interfacing with deep neural circuits typically requires the implantation of large-diameter rigid fibers, which can make this approach more invasive than its electrical counterpart^10,11^.

The ideal neural probe would combine optical and electrical modes while maintaining small cross-sectional dimensions and tunable lengths. The ability to bi-directionally interface with genetically defined neuron types and circuits is key to ultimately being able to understand how the nervous system computes and controls behavior. It is also fundamental for determining the mechanistic basis of sensorimotor disorders, defining how circuit activity is affected by injury, and how it might be restored or facilitated. Approaches to integrating optical and electrical modalities have ranged from adding fiber optics to existing Utah arrays^12^ to the Optetrode^13,14^ or other integrated electro-optical coaxial structures^15^. These technologies have shown great promise for simultaneous electrical recordings and optical stimulation *in* vivo. However, the need to reduce the device footprint to minimize immune responses for long-term recordings is still present^1,16–19^.

In this work, we present, to the best of our knowledge^1^, the smallest coaxial neural probe with a low impedance electrical channel surrounding a small central fiber optic core. These electro-optical mechanically flexible (EO-Flex) probes can be fabricated with diameters as small as 8 μm and lengths up to several millimeters using microfiber optic waveguide cores or even smaller diameters with nanofiber optic cores. They can be bonded directly to single-mode fibers (SMFs) to create detachable, low-loss optical interfaces that can be directly connected to standard optogenetic hardware. The EO-Flex probes’ simultaneous electrical recording and optical stimulation performance are demonstrated in the mouse brain. Our experiments show that the porous metal electrical channel provides excellent recording ability even with the probe’s small size. The low source-to-tip optical losses of < 10 dB allow robust optogenetic stimulation in transgenic or virally transduced mice expressing opsins in target cells. Implant studies show minimal immune responses, suggesting that the fully customizable probe and future high-density arrays should enable long-term interfacing with minimal disturbance to the surrounding neural tissue.

EO-Flex probes were fabricated using micro- and nanofiber optical cores (see Methods). Here, we will focus on mass-producible silica microfibers as the core that enables probes with lengths surpassing 3 mm while maintaining a diameter of < 12 μm (Supplementary Fig. 11a). However, the fabrication protocol is general and can be used with other optical cores, including subwavelength metal oxide nanofiber waveguides to produce ultra-miniaturized probes (Supplementary Fig. 3). In order to enable efficient coupling to optogenetic hardware, the microfibers were first placed on a silicon substrate, with one end of the fiber protruding the edge of the substrate, and then butt-coupled to a cleaved SMF (Fig. 1a). We used active alignment to maximize mode overlap between the microfiber and SMF. The coupling was locked in using a UV-curable optical adhesive droplet on the end of the SMF (Fig. 1b).

**Figure 1.**
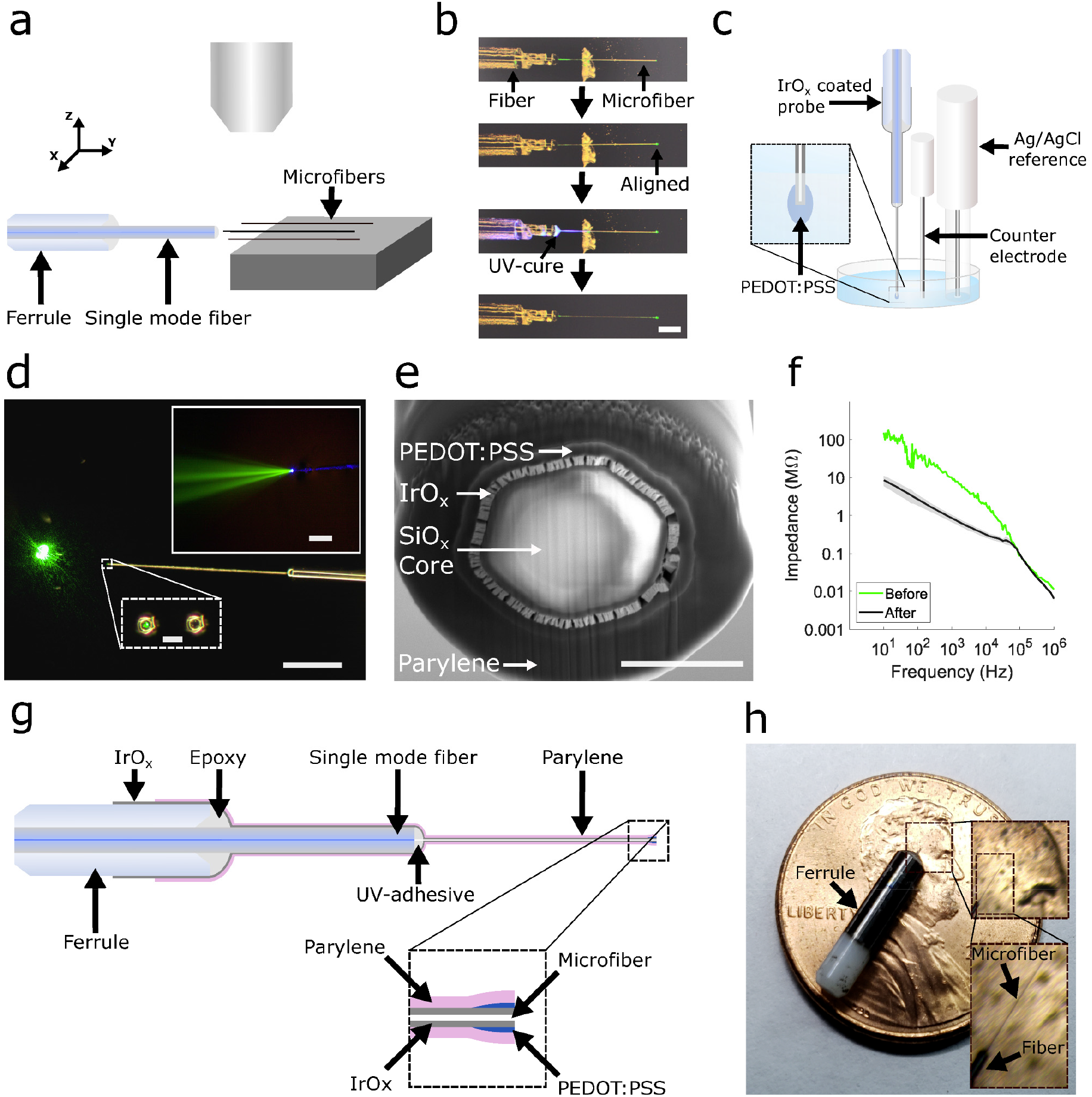
Fabrication of implantable EO-Flex probes along with optical and electrical characterization. (**a**) Silica microfibers of defined length are positioned on a silicon substrate to allow a single-mode fiber (SMF)-loaded ferrule to bond to the microfiber. (**b**) (from top to bottom) Photographs show the active alignment and bonding process of coupling the microfiber to the SMF. Scale bar 250 μm. (**c**) Schematic of the electrodeposition set-up for depositing PEDOT:PSS after the metal sputtering step (not shown). (**d**) Optical image of the light output of a probe from the side as the light reflects from a mirror, and from the cleaved end-facet (zoom-in insets) with and without laser light. (inset) Fluorescence image capturing the cone angle of the probe after submerging it in a dye solution and launching blue (442 nm) light into the probe. Scale bar 500 μm, 12 μm, and 60 μm respectively. (**e**) Cross-sectional electron micrograph of an EO-Flex probe after milling the end showing the exposed conductive rings along with the optical core. Scale bar 4 μm. (**f**) EIS data for milled probes with and without the PEDOT:PSS cladding. (the gray shaded area is one standard deviation; n=4 probes). (**g**) A cross-sectional view of the probe showing its various cladding layers. (**h**) Photograph of a completed EO-Flex probe with a zoom-in of the microfiber tip region. For scaling strategies, see Supplementary Fig. 11.

To create a robust detachable interface for in vivo testing, the SMF was inserted into a ceramic ferrule. The distal end of the ferrule assembly was machine polished to allow coupling to a patch cable (Fig. 1g-h). Other interface designs for different applications are conceivable (Supplementary Fig. 11).

To form a low noise conductive layer around the probe tip, a 379 ± 43 nm layer of iridium oxide (IrO_x_) was sputtered on the microfibers^20^ followed by a 362 ± 137 nm electrochemically deposited layer of poly(3,4-ethylene dioxythiophene) polystyrene sulfonate (PEDOT:PSS) (Fig. 1c)^20^. The porous nature of IrO_x_ allowed better adhesion of the conductive PEDOT:PSS layer and enhances the overall electrical performance of the probe. The probe was passivated with 1.76 ± 0.16 μm of Parylene-C to electrically isolate the probe and provide a biocompatible surface (Supplementary Fig. 2). In order to expose the electrical and optical surfaces, a focused ion beam was used to cleave off the tip (Fig. 1e; see Methods). Fig. 1g displays the final probe design and Fig. 1h shows a photograph of a completed probe. Through the combination of IrO_x_ and PEDOT:PSS, electrical impedances of < 1 MΩ at 1 kHz were achieved from electrode areas of < 15 μm^2^ (Fig. 1f).

The probes’ optical properties were first assessed by imaging the output cone angle in a dye solution (Fig. 1d, inset), which showed a uniform output with a divergence angle of 10 - 15°. Importantly, after the cladding layers are placed on the probe, no detectable scattering light is observed from the microfiber/SMF interface (Fig. 1d) as compared to the pre-cladding probe (Fig. 1b). The optical losses between a laser-coupled patch cable and the EO-Flex output were quantified using three different wavelengths (473 nm, 543 nm, 600 nm) with all devices showing < 7 dB (n=4). These values match up well with simulated results for an ~2 μm mode misalignment in the ferrule sleeve (Supplementary Fig. 1). Electrochemical impedance spectroscopy (EIS) was performed on the probes while submersed in a 1x phosphate-buffered solution (PBS). All probes fabricated and tested showed an average electrical impedance of 844 ± 179 kΩ at 1 kHz (n=4; Fig. 1f and Supplementary Fig. 2a-d) after cladding deposition and milling the tip, compared to >10 MΩ before PEDOT deposition.

To confirm that EO-Flex probes allow high-sensitivity electrical measurements in vivo, we performed simultaneous extracellular recordings and two-photon imaging in the cortex of isoflurane-anesthetized mice^21,22^. Imaging of fluorescently labeled cells (see Methods) enabled the monitoring of insertion and targeted movement of the probe through the tissue (Fig. 2a and Supplementary Fig. 8). Probes readily penetrated the exposed dura with minimal buckling when using water immersion and reached target regions in optically accessible cortical layer 2/3. When mounted to a three-axis micromanipulator, fine adjustment of the probe tip position, once inside the tissue, was feasible to optimize the signal-to-noise ratio and target individual neurons. However, the lateral movement was typically limited to < 30 μm. Using this approach, we acquired endogenous multi- and single-unit activity (Fig. 2b). Principle component analysis (PCA) and Gaussian clustering of electrical recordings were used to determine the number of distinct units (Fig. 2c). Spiking rates were calculated using a Bayesian Adaptive Kernel Smoother (BAKS) algorithm^23^ applied to the full duration of the recording (Fig. 2d). Fig. 2b-e shows a representative recording.

**Figure 2.**
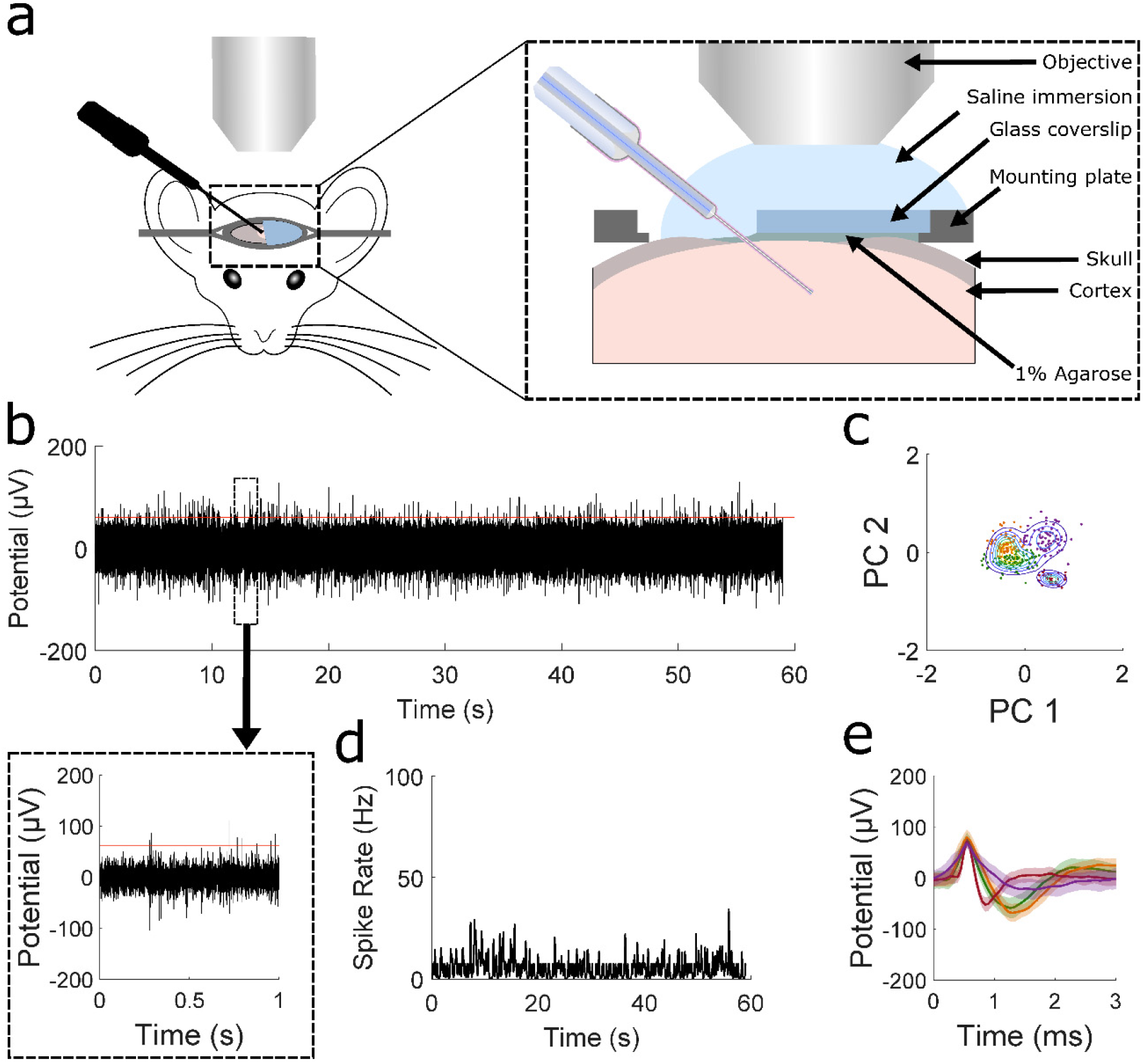
Extracellular neural recordings in the cortex of live mice using the EO-Flex probes. (**a**) Schematic showing the set-up used for visually guided electrical measurements. Two-photon imaging of the probe in relation to fluorescently labeled cells (see Methods) was used to verify and optimize the recording position. (inset) Zoom-in cross-sectional view of the surgical preparation for simultaneous imaging and electrical recordings. (**b**) Example EO-Flex recording showing spontaneous neural activity in cortical layer 2/3 (depth =250 μm) of an isoflurane-anesthetized mouse. The threshold (red line) defining a spike was set to 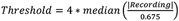 based on published literature^24^. (boxed region) A 1 s excerpt from the recording shows multi-unit activity. (**c**) PCA plot of the waveform clustering using established clustering methods^25^. (**d**) Spiking rate over the one-minute recording shown in (b) calculated using a Bayesian kernel estimation. (**e**) Average waveforms (solid lines) along with one standard deviation (shaded regions) for four clusters determined by PCA from the recording in (b).

Next, to demonstrate the EO-Flex probe’s ability to optically evoke neural activity while simultaneously electrically recording with the same probe, we performed experiments in anesthetized *Thy1-ChR2-YFP* mice with blue light-activated ion channel Channelrodopsin-2 (ChR2) expression in neurons. Probes were again inserted into cortical layer 2/3 under visual control. A 473 nm diode-pumped solid-state (DPSS) laser, suitable for exciting ChR2, was coupled into the probe, and stimulation parameters were swept systematically (Supplementary Figs. 4–7). We varied the stimulation frequency (10 Hz – 60 Hz), pulse width (0.6 - 9.8 ms), and output power (5 – 208 μW) to determine the optimal settings to excite ChR2-expressing neurons. Using waveform analysis on simultaneously recorded electrical activity, we found that a minimum power of 29 μW (2,849 mW mm^-2^) was required for firing of the cells in sync with the optical pulse train (Fig. S4). In the example recording shown in Fig. 3a-b, PCA combined with a mixed Gaussian fit for the clustering of the data yielded two primary clusters (Fig. 3c) with two different waveforms (Fig. 3d) occurring during the stimulation period (Fig. 3e-g). We found minimal interference (e.g., Becquerel effect) between the proximal optical and electrical channels, as demonstrated by retracting the EO-Flex probe away from ChR2-expressing neurons, or placing it in a buffer solution, while optically pumping at maximum power (208 μW) using the same optogenetic pulse trains (Supplementary Fig. 7). At this maximum power, neural circuits responded with minimal temporal lag (Fig. S4f) and could follow frequency stimuli up to 40 Hz before struggling to maintain synced firing (Supplementary Fig. 6).

**Figure 3.**
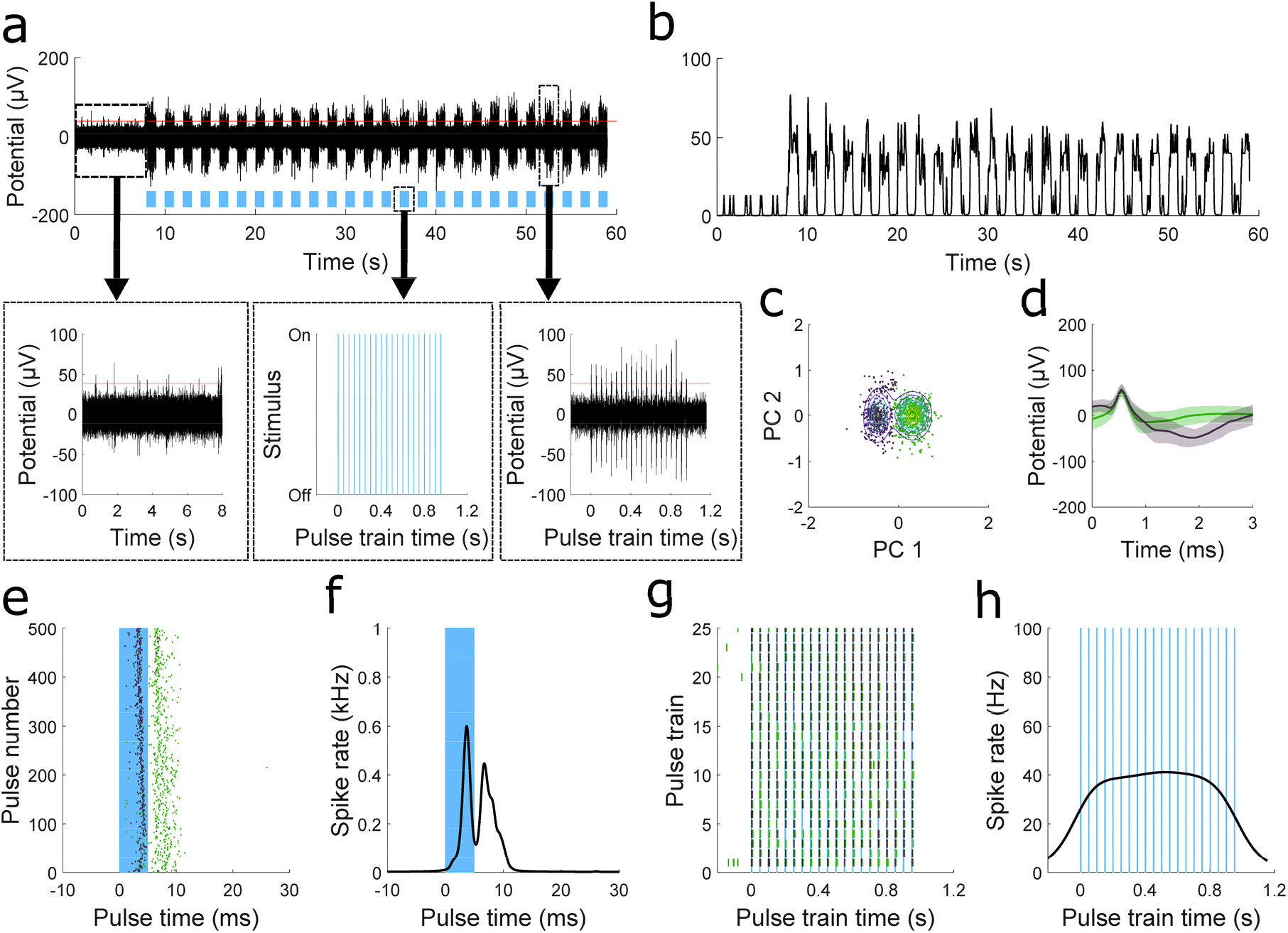
Concomitant optical stimulation and electrical recording with the EO-Flex probes in live *Thy1-ChR2-YFP* mice. (**a**) Optically evoked neural activity using a 20 Hz pulse train (blue bars) of 473 nm light (pulse width of 4.95 ms) at a tip power of 61 μW that is cycled on and off at 1 Hz. The threshold line (red) was set as defined^24^ in Fig. 2. (**b**) Spike rate plot for the recording in (a). (**c**) PCA plot for the optically evoked neural activity. (**d**) Average waveforms (solid line) for each cluster in (c) along with one standard deviation (shaded region). (**e**) Each cluster (color coordinated) plotted over time along with the window of a single pulse (blue bar). (**f**) Bayesian kernel smoothing estimate of spiking rate for each stimulation period. (**g**) Colored raster plot showing occurrence of waveforms from (d). (**h**) Calculated average spike rate over the 1 s duration of a single pulsing cycle.

The ability of EO-Flex probes to optically evoke neural activity was further verified by two-photon calcium imaging in *Vglut2-GCaMP6f* mice. Four to five weeks after the cortical injection of an AAV2-CaMKII-C1V1-mCherry vector (see Methods), expressing the green light-activated ion channel C1V1 in neurons, probes were inserted into layer 2/3 regions with C1V1 expression (Supplementary Fig. 8). A 556 nm DPSS laser was coupled to the EO-Flex probe, and stimulation parameters were swept while simultaneously monitoring neuronal calcium transients. Delivered optical pulses led to correlated calcium spiking in C1V1-positive neurons within the field of view (Supplementary Fig. 8). The successful optical evocation of neural activity was also verified by simultaneous electrical recordings (Supplementary Fig. 8e). Together, our in vivo data demonstrate the ability of EO-Flex probes to electrically record and optically modulate neural activity in the intact brain.

Lastly, we evaluated the brain’s response to probe implantation. EO-Flex probes were implanted into the cortex of *Cx3cr1^GFP/+^* mice with labeled microglia for seven days. A 250 μm-diameter multi-mode fiber, suitable for optogenetic experiments, was inserted using the same stereotaxic coordinates but on the opposite hemisphere for comparison. Serial brain sections were prepared that included both implantation sites. Tissue slices were co-stained with anti-GFAP and anti-NeuN antibodies to quantify reactive astrogliosis and neuronal loss, respectively (Supplementary Fig. 9–10). We found that implantation of the multi-mode fiber was associated with significant neuronal loss (Supplementary Fig. 10c), a 2.72 ± 0.35 fold increase in microglia numbers, and 2.62 ± 0.1 fold increase in GFAP levels (Supplementary Fig. 10d-e). In contrast, the EO-Flex probes showed no significant decrease in NeuN-positive cells or increase in microglia numbers and GFAP levels around the insertion site (Supplementary Fig. 10c-e). Together, these results indicate that tissue responses to EO-Flex probe insertion and potential animal behavior-related probe movement during the implantation period are negligible at a time point when inflammatory responses are typically most prominent^26^, and considerably smaller compared to standard probes used for optogenetic experiments.

EO-Flex probes allowed targeting and entraining of opsin-expressing cells at firing rates ranging from 10 Hz to 30 Hz (Supplementary Fig. 6). A minimum power threshold of 29 μW (2,849 mW mm^-2^ at the probe tip) was required for reliable activation of neurons, and while this irradiance is higher than in previous reports (1-10 mW mm^-2^)^14,27,28^, we did not see any optical degradation effects in the tissue (Supplementary Fig. 8). Monte Carlo simulations showed that, at 29 μW stimulation power, the optical power density drops below the optogenetic threshold of 1 mW mm^-2^ at around 1.2 mm from the tip. Even at the maximum stimulation power utilized in our optogenetic experiments (208 μW or 20,435 mW mm^-2^ at the probe tip, which drops to 2,168 mW mm^-2^ at a distance of 50 μm), we did not observe any adverse cellular effects (e.g., sustained changes in firing rate or calcium levels). Recent studies have suggested that continuous optical exposure with powers < 0.25 mW result in no temperature effects on neural activity (i.e., degraded electrical signals over the stimulation period)^29^. To ensure minimal optical heating effects on the neural tissue, we utilized short pulse widths and optical powers of less than 250 μW (Supplementary Fig. 4).

Developing probes that can reach deeper brain regions is straightforward with the developed fabrication protocols as virtually any microfiber length can be generated (Supplementary Fig. 11a). However, for a given set of cladding layers, the probe’s stiffness decreases with length. Therefore, longer probes might require additional tactics to overcome low buckling forces during the insertion process (e.g., dissolvable sugar coatings^30^, or rigid polymer layers). Alternatively, a surgical incision in the dura could facilitate probe insertion. Regardless of probe length, our implantation studies demonstrated that the small-footprint EO-Flex probes have a drastically reduced immune response in comparison to standard multi-mode fibers.

In summary, we report on the fabrication of novel multi-modal coaxial microprobes and demonstrate their ability to optically stimulate and electrically record from intrinsic neural circuits with minimal interference between the two modalities. The small footprint and high aspect ratio of the EO-Flex probes allow for minimally invasive interfacing with neural circuits. Further size reduction is possible with this coaxial design using smaller fiber optic cores, however, the tradeoff is an increase in optical losses and electrical impedance (Supplementary Fig. 3). Although the probes’ capabilities were only tested in the brain, as a platform with excellent control over probe diameter and length, the choice of cladding materials with various chemical compositions, inherent mechanically flexibility, and a clear route to scaling up probe densities (e.g., translating the cladding deposition process to fiber bundles) (Supplementary Fig. 11), this technology should find immediate applications as minimally invasive interfaces in diverse nervous system regions, including the spinal cord and peripheral nerves.

## Acknowledgements

We would like to thank Drs. Anis Husain and Rob Saperstein (Ziva Corporation) for technical discussions and electromagnetic simulations of early EO-Flex prototypes and designs. We would also like to thank Pavel Shekhtmeyster (Salk Institute) for technical assistance with the optogenetics experiments, and Ben Temple (Salk Institute) and Elischa Sanders (Salk Institute) for advice on the electrophysiological data analysis. This work was sponsored by the Defense Advanced Research Projects Agency (DARPA) Biological Technologies Office (BTO) Electrical Prescriptions (ElectRx) program under the auspices of Dr. Douglas Weber through the DARPA Contracts Management Office Grant/Contract No. HR0011-16-2-0027. This project was also supported by the UCSD Kavli Institute for Brain and Mind (Grant No. 2018-1492 to D.J.S. and A.N.) and the US National Institutes of Health (R01 NS108034, U19 NS112959, and U01 NS103522 to A.N., and P30CA014195 to the Salk Institute). This work was performed in part at the San Diego Nanotechnology Infrastructure (SDNI) of UCSD, a member of the National Nanotechnology Coordinated Infrastructure, which is supported by the National Science Foundation (Grant ECCS-1542148).

## Author Contributions

S.W., C.R., S.E., A.N. and D.J.S. conceptualized the probes; S.W., C.R., and J.N. fabricated, characterized, and tested the probes; S.W., E.C., and A.N. performed the biological experiments; S.E., A.N., and D.J.S. secured funding and supervised the study; S.W., A.N., and D.J.S. analyzed the data and wrote the paper; all authors reviewed and edited the manuscript.

## Supplementary Information

### Probe fabrication

EO-Flex probes were fabricated using one of two waveguides as the optical core: a) silica microfibers (SiO_x_) (Figs. 2 and Supplementary 2), or b) single crystalline tin dioxide (SnO_2_) nanofibers (Supplementary Fig. 3).

Silica microfibers (core and total diameters: 3.63 ± 0.31 μm and 5.60 ± 0.42 μm, respectively) with lengths varying between 500 μm and 1 cm were generated from leeched fiber optic bundles (Schott, Part no. 1573179). After cleaving, individual fibers were dispersed onto a silicon substrate. A tungsten needle mounted on a 3-axis micromanipulator was used to position the microfibers near a substrate edge with one end of the fiber being suspended > 100 μm from the edge.

The SnO_2_ nanofibers were synthesized using thermal evaporation of SnO powders at high temperatures according to published protocols^1^. Ceramic combustion boats were loaded with 1-5 grams of tin monoxide powder and placed in a tube furnace. The system was pumped down to < 1 mTorr as the furnace was turned on to 1000 °C. At operating temperature, system pressures were typically around 300 mTorr. The system was allowed to run for an hour, after which the furnace was turned off, and the system was allowed to cool while the vacuum pump remained on. The combustion boat was then removed, and nanowires found on the rim of the boat were transferred to a silicon substrate.

To enable efficient optical coupling of the waveguide to standard optogenetic hardware, a single-mode fiber (SMF) (Thorlabs S405-XP) with a mode field diameter (2.8-3.4 μm) slightly smaller than the microfiber core was chosen. This ensures high optical coupling efficiency. To create a robust detachable interface for in vivo testing, the SMF was inserted into a ceramic ferrule (Thorlabs CF126-10) and secured in place with quick cure epoxy (DevCon #20445). The ferrule assemblies were then machine polished until a smooth coupling interface was observed through a fiber inspection scope (Thorlabs, FS200), and the opposing fiber end (for coupling to the waveguide) was cleaved using a ruby scribe (Thorlabs S90R). The ferrule assembly was mounted on a three-axis stage, and the scribed end was maneuvered into a droplet of UV-cured optical adhesive (Norland Optical Adhesive 81) until a small droplet formed at the end. Efficient coupling between SMF and micro-/nanofiber was achieved using active alignment under an upright optical microscope (Nikon) equipped with a 0.4 NA 20x objective after coupling a 544 nm He-Ne laser source into the SMF. After maximizing power coupling into the waveguide by translating the SMF, the NOA 81 adhesive was secured by exposing it to UV light (325 nm line from a HeCd laser) for a duration of 30 s while continuously moving the beam around the droplet.

Before depositing the metal layer, the probe assemblies were placed in a custom aluminum block holder to mask the bottom part of the ferrule where light is coupled into the assembly. This ensured that the optical coupling interface was masked during metallization. This block was placed on a rotating plate inside a sputtering chamber (Denton Discovery 18). A thin (< 10 nm) adhesion layer of titanium (2.5 mTorr, 5 s, 200 Watts) was deposited, followed by a 300 nm thick layer of iridium oxide (IrO_x_) (12 mTorr, 15 min, 100 watts, 5 sccm O_2_ flow). Iridium oxide was chosen for its ~3x higher charge-injection capacity compared to conventional platinum layers^2^, and its porous nature, which increases the electrochemical surface area of the metal layer^2^.

Together, these procedures yielded ferrule assemblies for repeatable mounting to an in vivo imaging and optogenetics setup (Fig. 2). Alternative interface designs (e.g., for probe arrays) are shown in Supplementary Fig. 11.

### PEDOT-PSS deposition

To further lower the electrical impedance of the probes, a poly(3,4-ethylene dioxythiophene)-poly(styrene sulfonate) (PEDOT:PSS) layer was deposited on the IrO_x_. Probes were submersed (~ 100 μm of the probe tip) into a 0.01 M solution of EDOT (97%, Millipore-Sigma) with 2.5 mg/mL of poly(sodium styrene sulfonate) (PSS; Millipore-Sigma). The electrochemical deposition was performed using a platinum wire counter electrode and an Ag/AgCl reference electrode (CHI 111P, CH Instruments) connected to an electrochemical potentiostat (VersaSTAT4) operating in the galvanostatic mode set to run at a current of 200 nA for 5 - 30 s. This yielded a 362 ± 137 nm thick polymer layer (Supplementary Fig. 2a-d). The PEDOT-IrOx coated microfibers were then passivated with 1.5 – 2 μm of parylene-C using chemical vapor deposition (SCS Labcoater Deposition System; Specialty Coating Systems).

### In vitro probe characterization

A focused ion beam (FEI Scios Dualbeam) set to 5 nA at 30 kV was used to cleave off the end of the probe and expose the electrical and optical channels, revealing a final probe diameter of 8 - 12 μm for the microfiber cores. Electrochemical impedance spectroscopy (EIS) was carried out with the Versastat 4 to determine probe impedance in a 1x phosphate-buffered solution (PBS) using the same reference and counter electrodes described above. Optical coupling efficiency was determined by measuring light output from a fiber optic patch cable (Thorlabs, P1405B-FC-5) using three light sources (473 nm, 543 nm, and 673 nm) interchangeably coupled into the cable. Light power was measured by placing the ferrule 5-10 mm away from the detector head of a digital power meter (Thorlabs, PM100D). A ceramic ferrule sleeve (Thorlabs, ADAL1) was then slid halfway onto the patch cable, and different EO-Flex probes were slid into the opposite end to couple light through. Light power from the tip of the EO-Flex probes was measured using a similar protocol to the patch cable.

### Animal subjects

All live animal procedures were performed following the guidelines of the National Institutes of Health (NIH) and were approved by the Institutional Animal Care and Use Committee (IACUC) at the Salk Institute under protocol number 13-00022. For combined optogenetic and electrophysiological experiments, we used *Thy1-ChR2-YFP* mice (stock #007612; Jackson Laboratories); for combined calcium imaging, optogenetics, and electrophysiological experiments we used AAV2-CaMKII-C1V1-mCherry-injected *Vglut2-GCaMP6f* mice (a custom cross between *Vglut2-Cre* knock-in and *Ai95D* mice; stock #028863 and #024105, respectively; Jackson Laboratories); for immune response studies we used *Cx3cr1^GFP/+^* mice (stock #005582; Jackson Laboratories).

### Stereotactic viral vector injection

Surgical procedures closely followed previously established protocols^3,4^. Briefly, thin-wall glass pipettes were pulled on a Sutter Flaming/Brown micropipette puller (model P-97). Pipette tips were cut at an acute angle under 10x magnification using sterile techniques. Tip diameters were typically 15–20 μm. Pipettes that did not result in sharp bevels nor had larger tip diameters were discarded. Millimeter tick marks were made on each pulled needle to measure the virus volume injected into the brain.

Mice were anesthetized with isoflurane (4% for induction; 1%-1.5% for maintenance) and positioned in a computer-assisted stereotactic system with digital coordinate readout and atlas targeting (Angle Two, Leica). Body temperature was maintained at 36°C–37°C with a DC temperature controller, and ophthalmic ointment was used to prevent eyes from drying. A small amount of depilator cream (Nair) was used to remove hair at the designated skin incision site. The skin was cleaned and sterilized with a two-stage scrub of betadine and 70% ethanol. A midline incision was made beginning just posterior to the eyes, and ending just passed the lambda suture. The scalp was pulled open, and periosteum cleaned using scalpel and forceps to expose the desired hemisphere for calibrating the digital atlas and coordinate marking. Once reference points (bregma and lambda) were positioned using the pipette tip and entered into the program, the desired target was set on the digital atlas. The injection pipette was carefully moved to the target site (coordinates: AP −1.5 mm, ML 1.5 mm). Next, the craniotomy site was marked, and an electrical micro-drill with a fluted bit (0.5 mm tip diameter) was used to thin a 0.5–1 mm diameter part of the bone over the target injection site. Once the bone was thin enough to flex gently, a sterile 30G needle with an attached syringe was used to carefully cut and lift a small (0.3–0.4 mm) segment of bone.

For injection, a drop of the virus was carefully pipetted onto parafilm (1–2 μl) for filling the pulled injection needle with the desired volume. Once loaded with sufficient volume, the injection needle was slowly lowered into the brain until the target depth (DV 0.2 mm) was reached. Manual pressure was applied using a 30-ml syringe connected by shrink tubing, and 0.4 μl of the AAV2-CaMKII-C1V1-mCherry vector (6.1E+12 VP/ml; undiluted; UNC Vector Core) was slowly injected over 5–10 min. Once the virus was injected, the syringe’s pressure valve was locked. The position was maintained for approximately 10 min to allow the virus to spread and to avoid backflow upon needle retraction. Following the injection, the skin was sutured along the incision. Mice were given subcutaneous Buprenex SR (0.5 mg per kg) and allowed to recover before placement in their home cage. The vector was allowed to express for 4-5 weeks before in vivo recordings.

### Animal preparation for in vivo recordings

Surgical procedures closely followed established protocols^5,6^. Briefly, mice were anesthetized with isoflurane (4-5% for induction; 1%-1.5% for maintenance) and implanted with a head plate on a custom surgical bed (Thorlabs). Body temperature was maintained at 36°C–37°C with a DC temperature control system, and ophthalmic ointment was used to prevent the eyes from drying. Depilator cream (Nair) was used to remove hair at the designated skin incision site. The skin was thoroughly cleansed and disinfected with a two-stage scrub of betadine and 70% ethanol. A scalp portion was surgically removed to expose frontal, parietal, and interparietal skull segments. Scalp edges were attached to the lateral sides of the skull using a tissue-compatible adhesive (Vetbond; 3M). A custom-machined metal plate was affixed to the skull with dental cement (cat. #H00335; Coltene Whaledent), allowing the head to be stabilized with a custom holder. An approximately 2 mm x 4 mm diameter craniotomy was made over the target area (e.g., AAV injection site). The dura mater overlying the cortex was kept intact. A 1% agarose solution and coverslip were applied to the exposed cortical tissue. To facilitate probe entry into the tissue, the agarose and coverslip were cut on one side to be flush with the craniotomy, allowing direct cortical access through the agarose. The coverslip was affixed to the skull with dental cement to control tissue motion. Recordings commenced immediately after optical window preparation. The depth of anesthesia was monitored throughout the experiment and adjusted as needed to maintain a breath rate of approximately 55-65 breaths per minute. Saline was supplemented as needed to compensate for fluid loss.

### In vivo electrophysiology

To characterize the electrophysiological properties of the EO-Flex probes, we performed extracellular single- and multi-unit recordings in the cortex of isoflurane-anesthetized mice, as previously described^5,7^. The EO-Flex probes’ electrical channel was connected to the positive terminal of a high impedance head stage (Model 1800 microelectrode AC amplifier, A-M Systems) with the negative terminal and ground attached to an Ag/AgCl wire inserted above the cerebellar cortex. To allow targeted tissue insertion and precise positioning, the probe was mounted to a motorized micromanipulator (MP-225, Sutter Instrument Company) angled at approximately 30 degrees relative to the optical axis of the microscope. After positioning the tip of the EO-Flex probe near the edge of the craniotomy, a few drops of physiological saline were pipetted onto the exposed agarose/cortex interface to facilitate mechanical insertion through the agarose and dura (Fig. 2a). The probe tip was positioned near neuronal cell bodies in upper cortical layers (mostly layer 2/3) of fluorescent indicator-expressing transgenic mice using two-photon imaging. Precise positioning was aided by passing the output from the differential amplifier through a speaker to serve as auditory feedback for probe proximity to active cells. The raw electrode signal was amplified, filtered (low cut-off, 300 Hz; high cut-off, 5 kHz; gain, 1000x), digitized (20 kHz), and stored on disk for offline analysis.

### Two-photon microscopy

In vivo imaging was performed as previously described^3,5^. Briefly, a Movable Objective Microscope (Sutter Instrument Company) equipped with a pulsed femtosecond Ti:Sapphire laser (Chameleon Ultra II, Coherent) with two fluorescence detection channels was used for imaging (emission filters: ET525/70M and ET605/70M (Chroma); dichroic beam splitter: 565DCXR (Chroma); photomultiplier tubes: H7422-40 GaAsP (Hamamatsu)). The laser excitation wavelength was set to 920 nm. The average laser power was <10-15 mW at the tissue surface and adjusted with depth to compensate for signal loss due to scattering and absorption. A 16x 0.8 NA (CFI75, Nikon) or 40x 0.8 NA (LUMPLFLN, Olympus) water-immersion objective was used for light delivery and collection. Spontaneous and optically evoked calcium activity was recorded in optical planes near the probe tip (frame rate, 8.14 Hz). To minimize the Becquerel effect mediated artifacts in electrical recordings, the imaging laser power was kept to a minimum. To record optically evoked calcium transients in optogenetic experiments, we synchronized the image frame rate with optical pulse train delivery and adjusted the phase such that regions in front of the probe tip were scanned when the DPSS laser was off (Supplementary Fig. 8).

### In vivo optogenetics

To excite ChR2 or C1V1, respectively, the light from a 200-mW 473 nm or 556 nm DPSS laser (CNI), directly modulated by an external function generator signal, was coupled into the probe. Light coupling into the probe was achieved by sliding the polished end of the ferrule into a ceramic sleeve and then sliding it onto the end of a custom fiber patch cable. Each stimulation trial lasted around 60 s, with the initial 5-10 s designated to recording spontaneous activity before the optical pulse train was delivered (stimulation power, 6-208 μW; pulse width, 0.6-9.8 ms; stimulation frequency, 10 - 50 Hz; duration, 1s; inter-stimulus-interval, 1s between pulse trains).

### Tissue response assessment

*Cx3cr1^GFP/+^* mice with labeled microglia were implanted on a given hemisphere ~1.45 mm from midline with an EO-Flex probe or a 250 μm diameter multimode fiber suited for optogenetic deep brain stimulation. For implantation, an electrical micro-drill with a fluted bit (0.5 mm tip diameter) was used to thin a 0.5–1 mm diameter part of the bone. Once the bone was thin enough to flex gently, a sterile 30G needle with an attached syringe was used to carefully cut and lift a small (0.3–0.4 mm) segment of bone. The probe or multimode fiber was advanced through this opening under visual control to a depth of approximately 1 mm using a computer-assisted stereotactic system (Angle Two, Leica). Dental cement was used to secure the devices in place. The firm bonding of the dental cement to the skull was facilitated by scarifying it with a bone scraper (Fine Science Tools). To distinguish surgery from probe related tissue responses, we performed additional craniotomies of the same size 0.7 mm lateral to the device implantation site (Supplementary Fig. 10). To assess tissue inflammatory responses, mice were sacrificed 6-7 days after device implantation using CO_2_ asphyxiation following IACUC guidelines. Transcardial perfusion was performed with 10% sucrose in phosphate-buffered saline (PBS), followed by freshly prepared 4% PFA in PBS. Both hemispheres were post-fixed in 4% PFA in PBS overnight and subsequently infiltrated in 30% sucrose in PBS for one day and flash frozen in TBS tissue freezing medium. The implanted hemispheres were coronally cryo-sectioned at 20 μm, air-dried overnight, and subsequently processed for staining. Sections were incubated overnight at 4°C with primary antibody diluted in blocking buffer, then washed in PBS 0.1% Tween-20, and incubated for two hours at 22-24 °C in the dark with fluorophore-coupled secondary antibodies. Sections were washed, sealed with Prolong Gold Antifade Mountant (Thermo Fisher Scientific), and stored at 4 °C. Primary antibodies included anti-GFAP (mouse monoclonal; EMD Millipore; cat. #MAB3402; RRID: AB_94844; 1:250 dilution) and anti-NeuN (rabbit polyclonal; EMD Millipore; cat. #ABN78; RRID: AB_10807945; 1:100 dilution). Secondary antibodies (1:100) included Alexa Fluor 405 goat antirabbit (Thermo Fisher Scientific; cat. #A-31556; RRID: AB_221605) and Alexa Fluor 633 goat anti-mouse (Thermo Fisher Scientific; cat. #A-21052; RRID: AB_2535719). Confocal imaging of stained tissue sections was performed on a Zeiss LSM 710. Three-channel tiled z-stacks were acquired to produce images of whole-tissue sections (Supplementary Fig. 9). Image size was 1,024 x 1,024 pixels stitched into 3-5 x 3-5 tiles. Images were taken with an Olympus 20x 0.8 NA air-matched objective.

### Data processing and statistical analyses

Neural activity was considered a spike if its amplitude crossed a threshold^8^ determined by 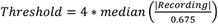. All observed spikes were then sorted according to the first two principal components into clusters using a mixed Gaussian fitting with the number of clusters optimized according to the Calinksi-Harabasz metric for cluster analysis^9^. Monte Carlo simulations were used to determine the propagation and illumination volume of the EO-Flex probe at different powers (Supplementary Fig. 4b).

Peristimulus plots correlating optical stimuli with spiking events were calculated using kernel bandwidth optimization, which has been shown to accurately estimate the underlying spiking rate^4^ (Fig. 3b). For stimulation frequencies of 10, 20, and 30 Hz, we saw estimated firing rates closely following input frequencies when accounting for the echo after stimulation. In comparison, frequencies of 40 and 50 Hz initially tracked the input frequency for the first couple of light pulses, but eventually, the firing rate decreased as the optical train progressed (Supplementary Fig. 6g).

Analysis of the two-photon calcium imaging data was performed using Suite2p^10^ (Supplementary Fig. 8). Optically evoked calcium spiking was observed in optical planes near the probe tip. Our analysis focused on the optical planes in which at least three cellular-size regions of interest (ROIs) consistently responded throughout the stimulation period (Supplementary Fig. 8).

Immunostaining data were processed, analyzed, and plotted using ImageJ, Imaris, and Prism software. All data are represented as mean ± s.e.m. Group sample sizes were chosen based on previous studies and power analysis. The following convention was used to indicate P values: ‘ns’ indicates P>0.05, ‘*’ indicates 0.01<P≤0.05, ‘**’ indicates 0.001<P≤0.01, and ‘***’ indicates 0.0001<P≤0.001.

**Supplemental Figure S1.**
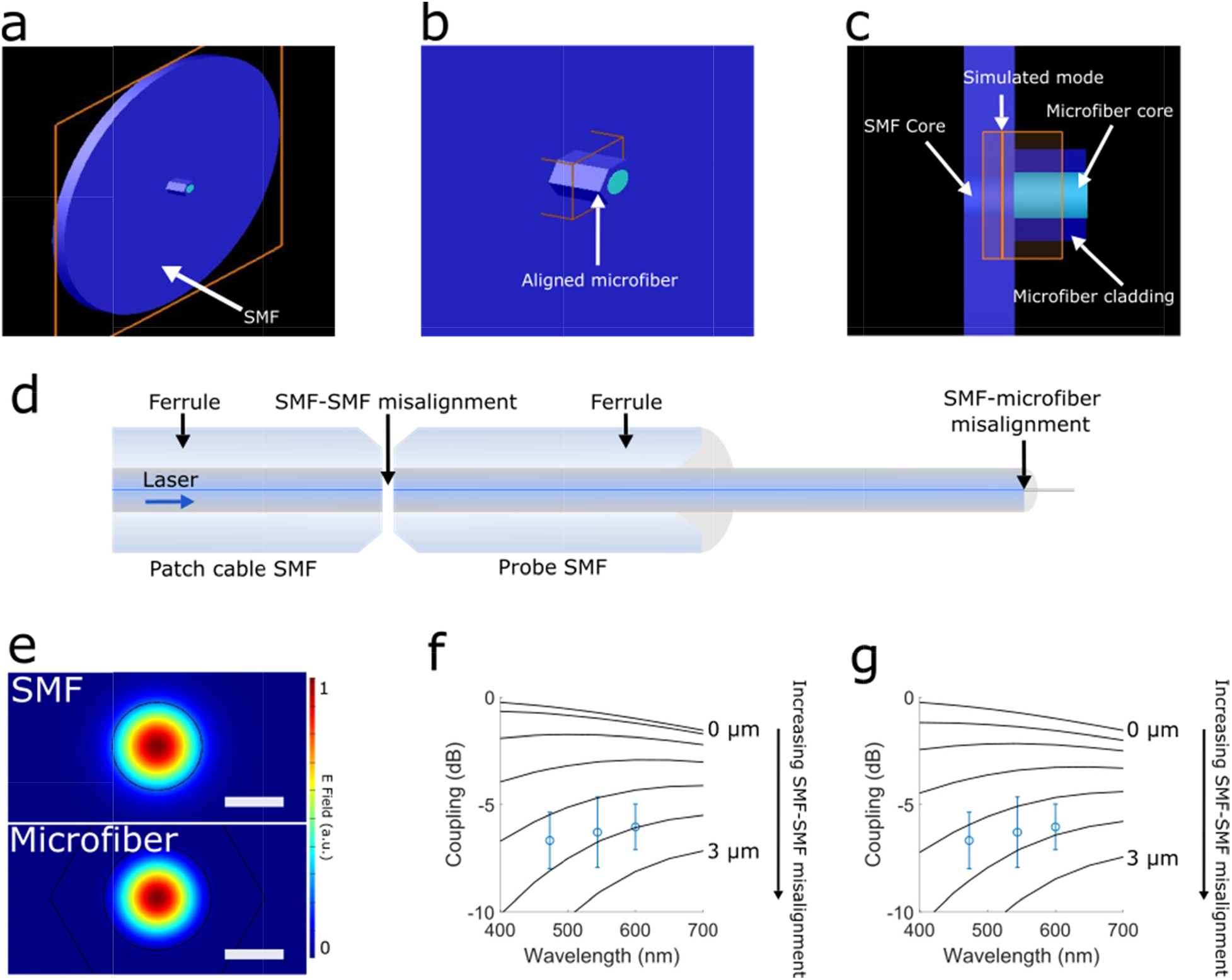
Finite element modelling of EM modes (Lumerical MODE) to investigate the theoretical optical coupling between the single-mode fiber (SMF) and microfiber. (**a**) Simulation geometry of a SMF fiber that has its core perfectly aligned with the core of the microfiber. (**b**) Zoom-in of the SMF-microfiber interface showing a 10 nm mesh used to quantify the coupling efficiency. (**c**) Side profile of (b) with design and labels of the model. (**d**) Schematic of the optical coupling into the probe from the laser source showing where the mode misalignment was simulated (SMF-SMF and SMF-microfiber). (**e**) Mode profile simulated at 473 nm for both the SMF and the microfiber. Scale bars 2 μm. (**f**) Simulated coupling efficiency assuming no misalignment between the SMF-microfiber interface and misalignment of the SMF-SMF interface from 0 to 3 μm (solid lines indicate every 0.5 μm of misalignment). Average coupling of 4 probes measured at 473nm, 543nm, and 600nm are overlaid on simulated curves. (**g**) Calculated coupling efficiency assuming the SMF-microfiber interface has 500 nm of misalignment and similar misalignment of the SMF-SMF interface as in (f). The same measured data set in (f) is overlaid on the new coupling curves. Comparison of simulation and measurements shows that optical coupling losses are mostly due to SMF-SMF misalignment (measured losses fall between 2 - 2.5 μm of SMF-SMF misalignment).

**Supplemental Figure S2.**
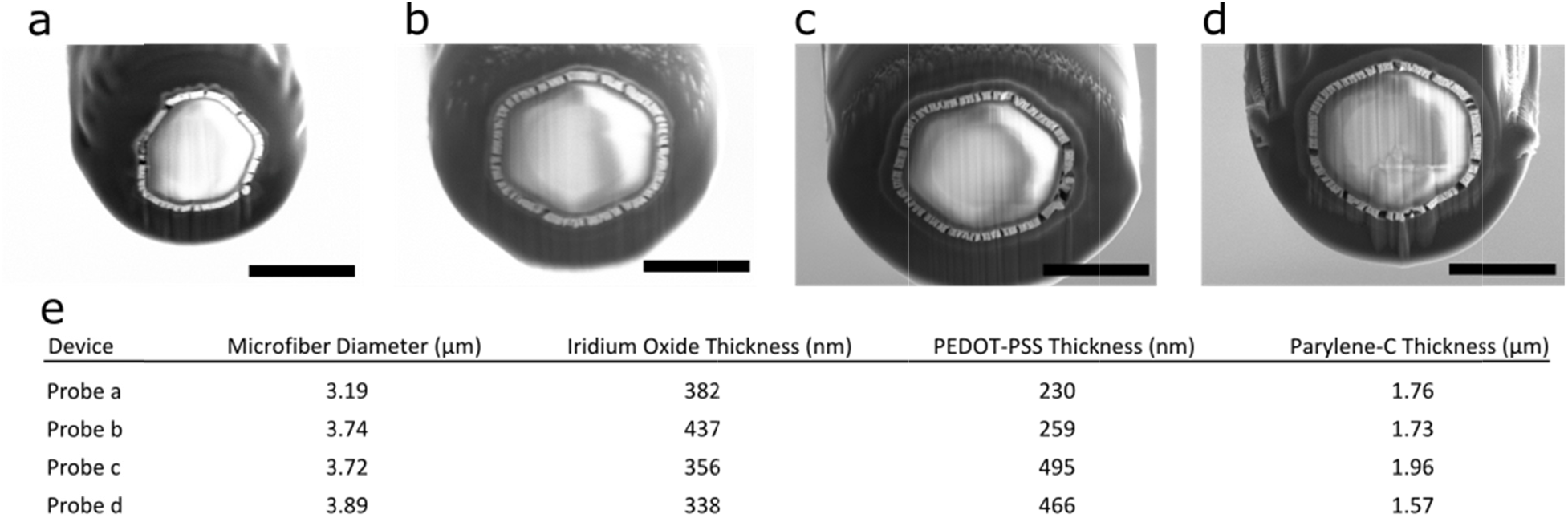
(**a**) - (**d**) Electron micrographs of four microfiber EO-Flex probes with different PEDOT:PSS deposition conditions. Data was used to optimize the polymer thickness and quantify the dimensions of the other cladding layer. Scale bars are 5 μm (a) and 4 μm (b-d). (**e**) Table of the different thicknesses for each layer deposited on the probes.

**Supplemental Figure S3.**
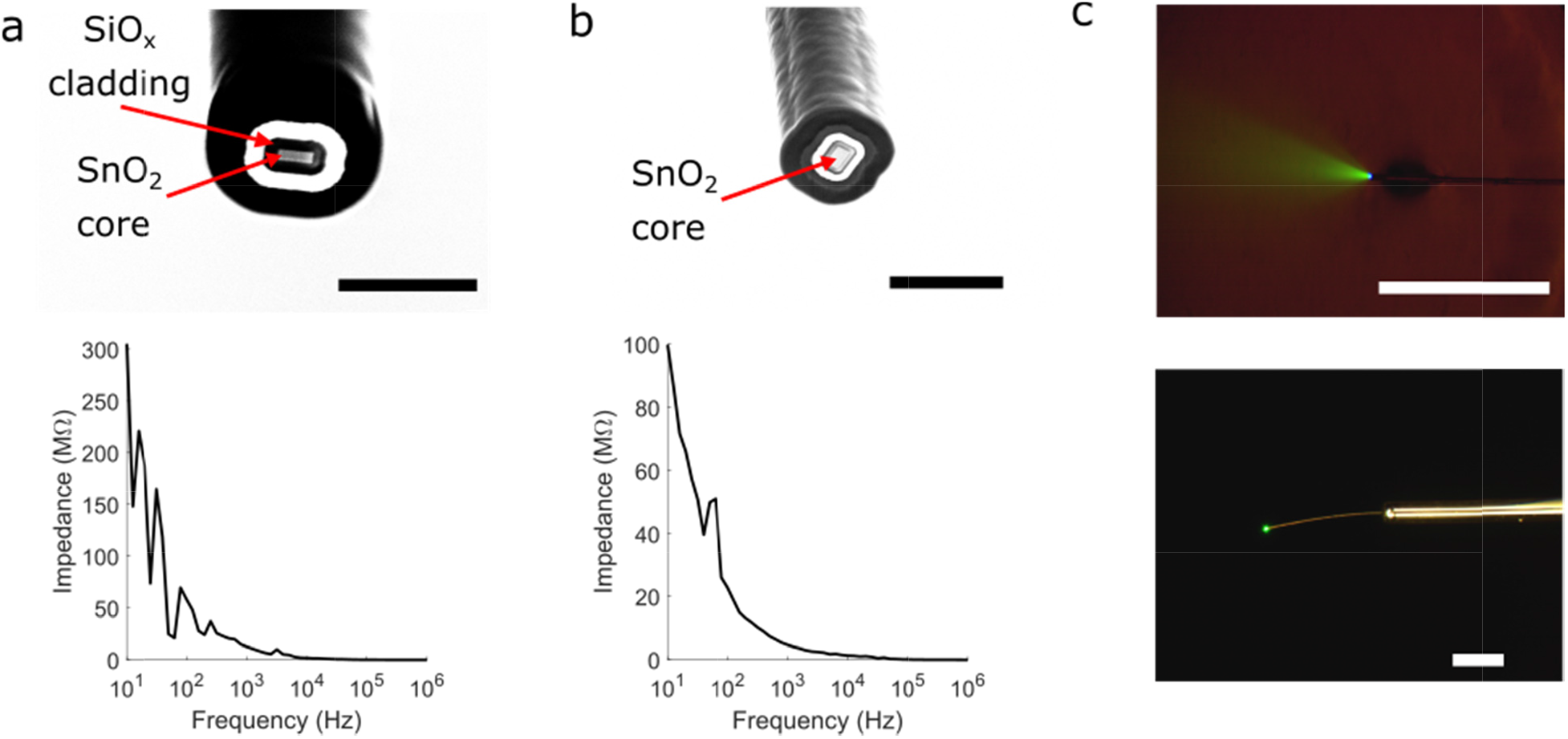
EO-Flex probes fabricated with a single crystalline tin dioxide (SnO_2_) nanofiber waveguide as the optical core. (**a**) (top) Electron micrograph of a SnO_2_ EO-Flex probe fabricated without the PEDOT-PSS layer. (bottom) EIS data of the probe showing an impedance of >20 MΩ at 1kHz. Scale bar 5 μm. (**b**) (top) Electron micrograph of a SnO_2_ EO-Flex probe fabricated with the PEDOT-PSS layer. (bottom) EIS data of the probe showing a significant reduction in the impedance down to 5 MΩ at 1 kHz. Scale bar 10 μm. (**c**) (top) The optical output of a SnO_2_ EO-Flex probe in a fluorescent dye solution showing the exiting cone angle. Scale bar 250 μm. (bottom) Optical image of the freestanding probe showing no light scattering at the SnO_2_-SMF interface after cladding deposition. Scale bar 250 μm.

**Supplemental Figure S4.**
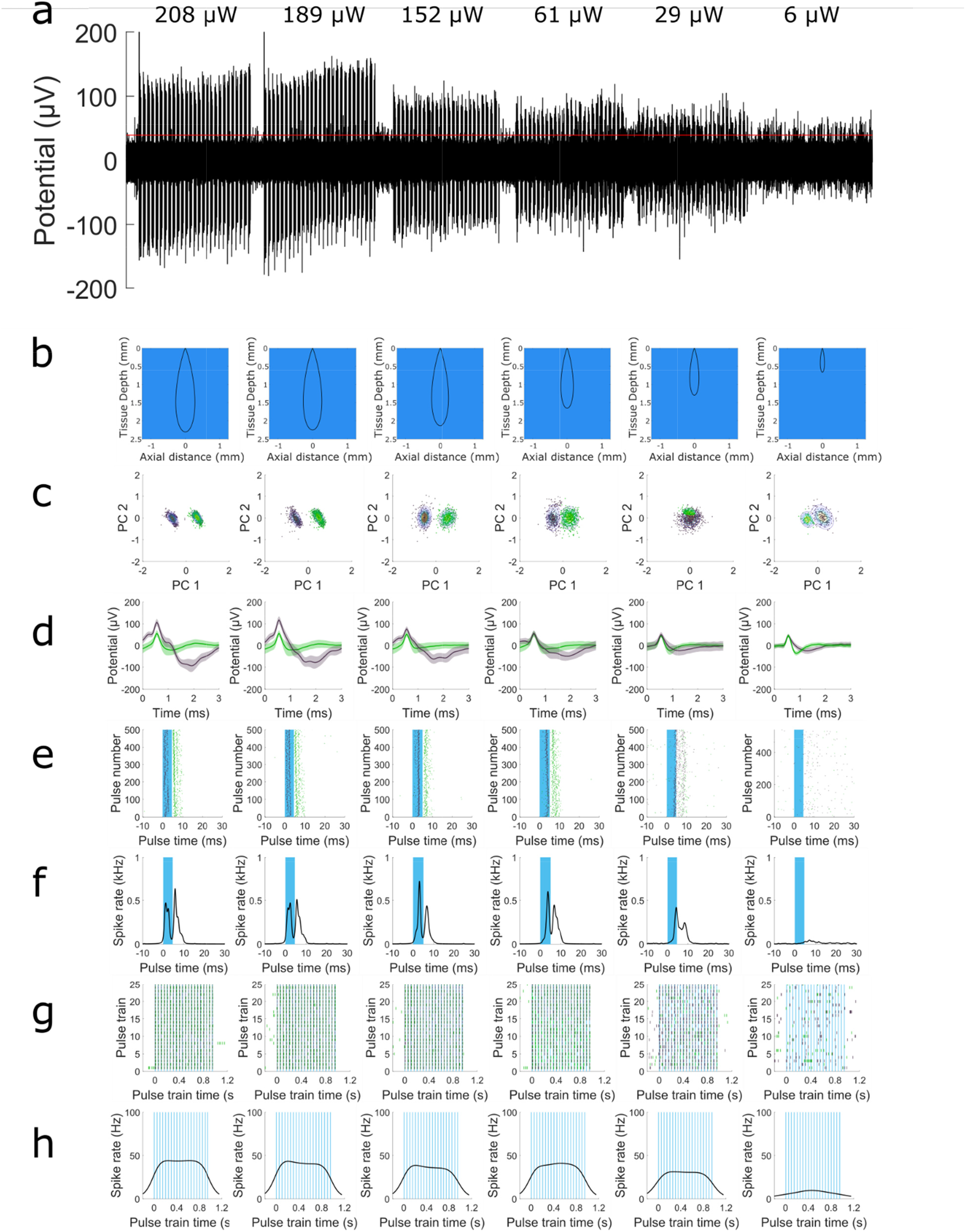
EO-Flex testing in layer 2/3 of a live *Thy1-ChR2-YFP* mouse as a function of optical stimulation power. All other stimulation parameters were held constant (pulse width, 4.5 ms; stimulation frequency, 20 Hz; on/off cycling, 1 Hz). Recording depth was ~250 μm. (**a**) Optically evoked neural activity using EO-Flex output powers ranging from 208 μW (18,832 mW mm^-2^) to 6 μW (543 mW mm^-2^). (**b**) Monte Carlo simulations of the scattering and absorption in neural tissue for each of the power values in (a). The solid line indicates where irradiance has fallen to 1 mW mm^-2^. The simulations showed that a light intensity of 208 μW could propagate up to 2.4 mm from the probe tip before the irradiance dropped below 1 mW mm^-2^. Simulation parameters were taken from recent studies^11^ which estimated scattering and absorption coefficients of 0.125 mm^-1^ and 7.37 mm^-1^, respectively. (**c**) The first two principal components (PCs) of respective electrical recordings plotted with the Calinski-Harabasz metric for determination of the number of clusters for mixed Gaussian fitting. (**d**) The average waveform for each cluster (solid line) from (c) with the shaded region representing one standard deviation. (**e**) Peri-stimulus plots for all optical pulses with spikes color-coordinated with the cluster from which they come. (**f**) Bayesian adaptive kernel smoother (BAKS) estimation for the firing rate over the time window around the optical pulses. (**g**) Peri-stimulus plot for each optical pulse train from high to low power. (**h**) BAKS estimation for the firing rate over the pulse train window.

**Supplemental Figure S5.**
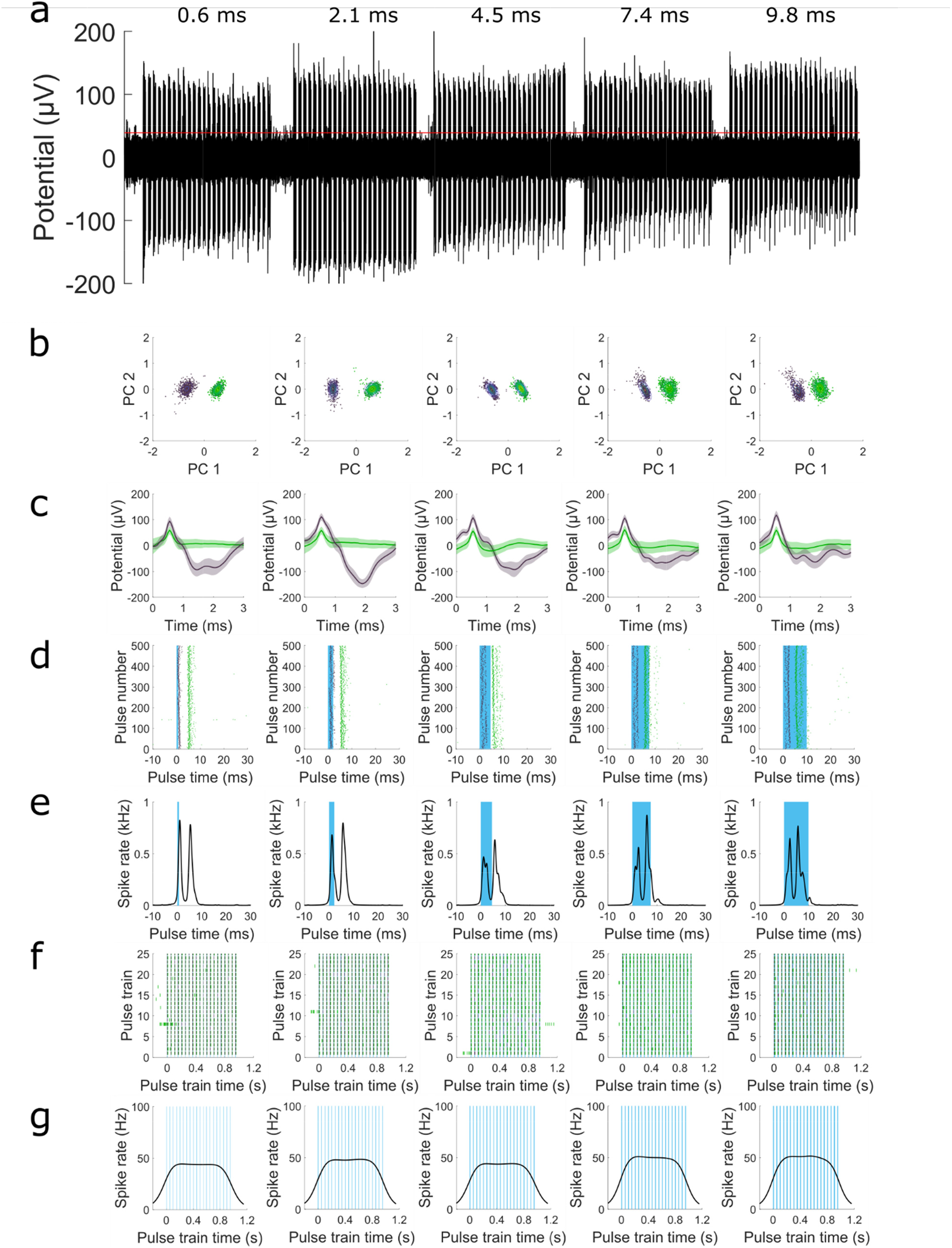
EO-Flex testing in layer 2/3 of a live *Thy1-ChR2-YFP* mouse as a function of the optical pulse width. All other stimulation parameters were held constant (optical stimulation power, 208 μW; stimulation frequency, 20 Hz; on/off cycling, 1 Hz). Recording depth was ~250 μm. (**a**) Optically evoked neural activity using pulse widths ranging from 0.6 ms to 9.8 ms. (**b**) The first two principal components (PCs) of respective electrical recordings plotted with the Calinski-Harabasz metric for determining the number of clusters for mixed Gaussian fitting. (**c**) Average waveform (solid line) for each cluster from (b) with the shaded region representing one standard deviation. (**d**) Peri-stimulus plots for all optical pulses with spikes color-coordinated with the cluster from which they come. (**e**) Bayesian adaptive kernel smoother (BAKS) estimation of firing rate over the time window around the optical pulses. (**f**) Peri-stimulus plot for each optical pulse train as a function of the optical pulse width. (**g**) BAKS estimation for the firing rate over the pulse train window.

**Supplemental Figure S6.**
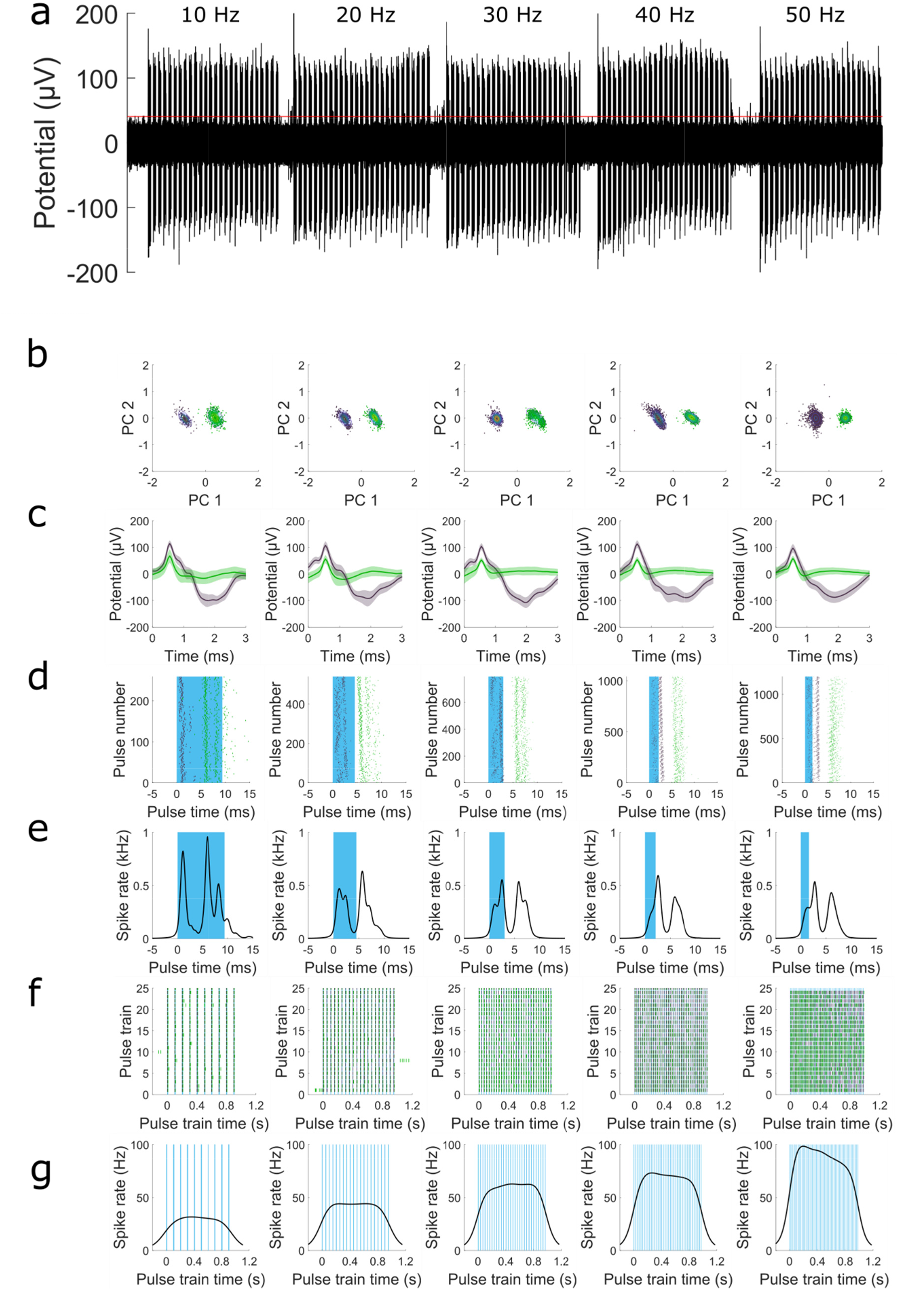
EO-Flex testing in layer 2/3 of a live *Thy1-ChR2-YFP* mouse as a function of stimulation frequency. All other stimulation parameters were held constant (optical stimulation power, 208 μW; duty cycle, 10%; on/off cycling, 1 Hz). Recording depth was ~250 μm. (**a**) Optically evoked neural activity using stimulation frequencies ranging from 10 Hz to 50 Hz. (**b**) The first two principal components (PCs) of respective electrical recordings plotted with the Calinski-Harabasz metric for determining the number of clusters for mixed Gaussian fitting. (**c**) Average waveform (solid line) for each cluster from (b) with the shaded region representing one standard deviation. (**d**) Peri-stimulus plots for all optical pulses with spikes color-coordinated with the cluster from which they come. (**e**) Bayesian adaptive kernel smoother (BAKS) estimation of the firing rate over the short time window around the optical pulses. (**f**) Peri-stimulus plot for each optical pulse as a function of the stimulation frequency. (**g**) BAKS estimation for the firing rate over the pulse train window.

**Supplemental Figure S7.**
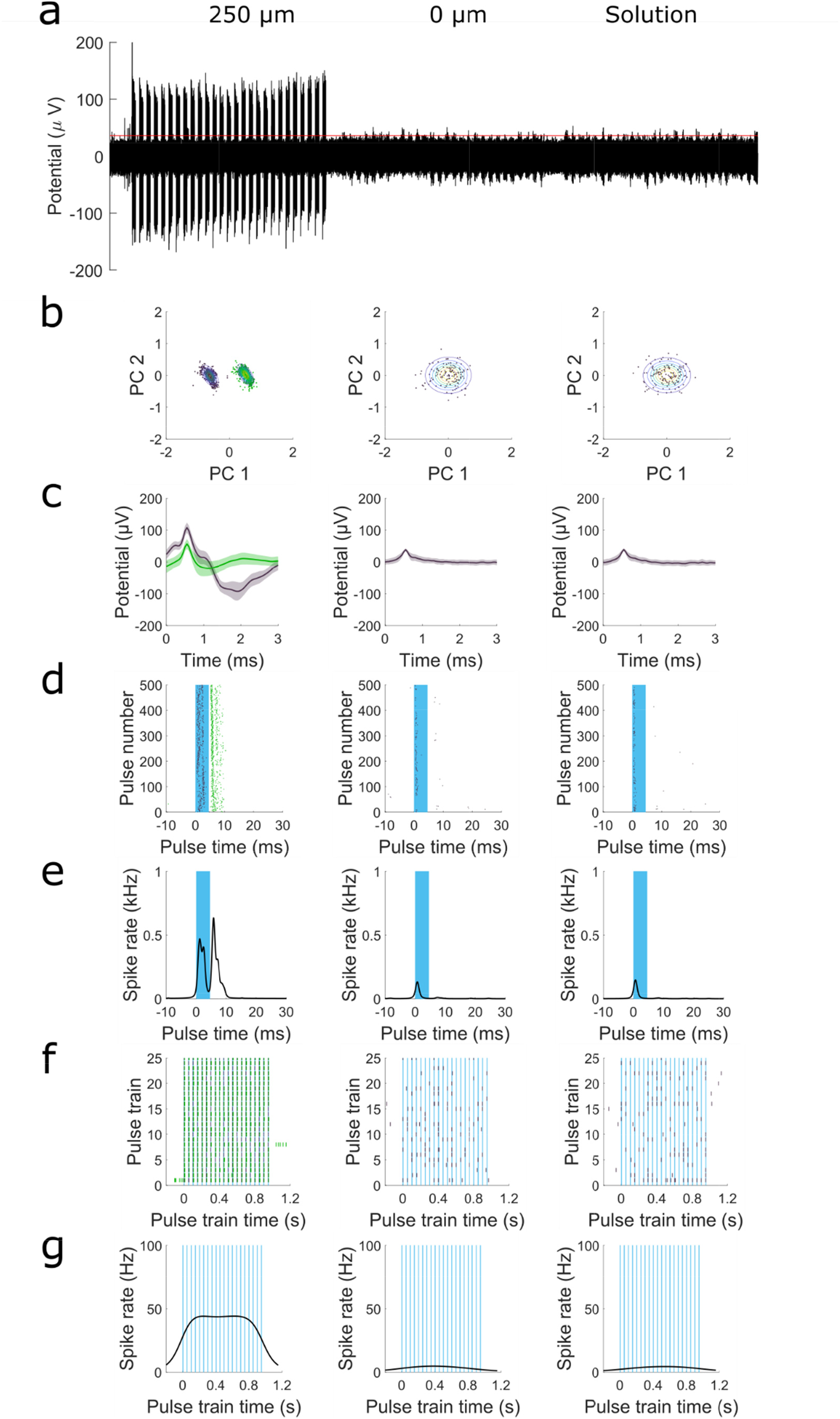
EO-Flex testing in a live *Thy1-ChR2-YFP* mouse as a function of recording depth. All other stimulation parameters were held constant (optical stimulation power, 208 μW; stimulation frequency, 20 Hz; pulse width, 4.5 ms; on/off cycling, 1 Hz). Recording depths were ~250 μm, 0 μm (i.e., at the agarose/brain interface), and in the saline solution above the craniotomy. (**a**) Optically evoked activity at different depths. (**b**) The first two principal components (PCs) of respective electrical recordings plotted with the Calinski-Harabasz metric for determining the number of clusters for mixed Gaussian fitting. (**c**) Average waveform (solid line) for each cluster from (b) with the shaded region representing one standard deviation. (**d**) Peri-stimulus plots for all optical pulses with spikes color-coordinated with the cluster from which they come. (**e**) Bayesian adaptive kernel smoother (BAKS) estimation of the firing rate over the short time window around the optical pulses. (**f**) Peri-stimulus plot for each optical pulse train for the different recording depths. (**g**) BAKS estimation for the firing rate over the pulse train window.

**Supplemental Figure S8.**
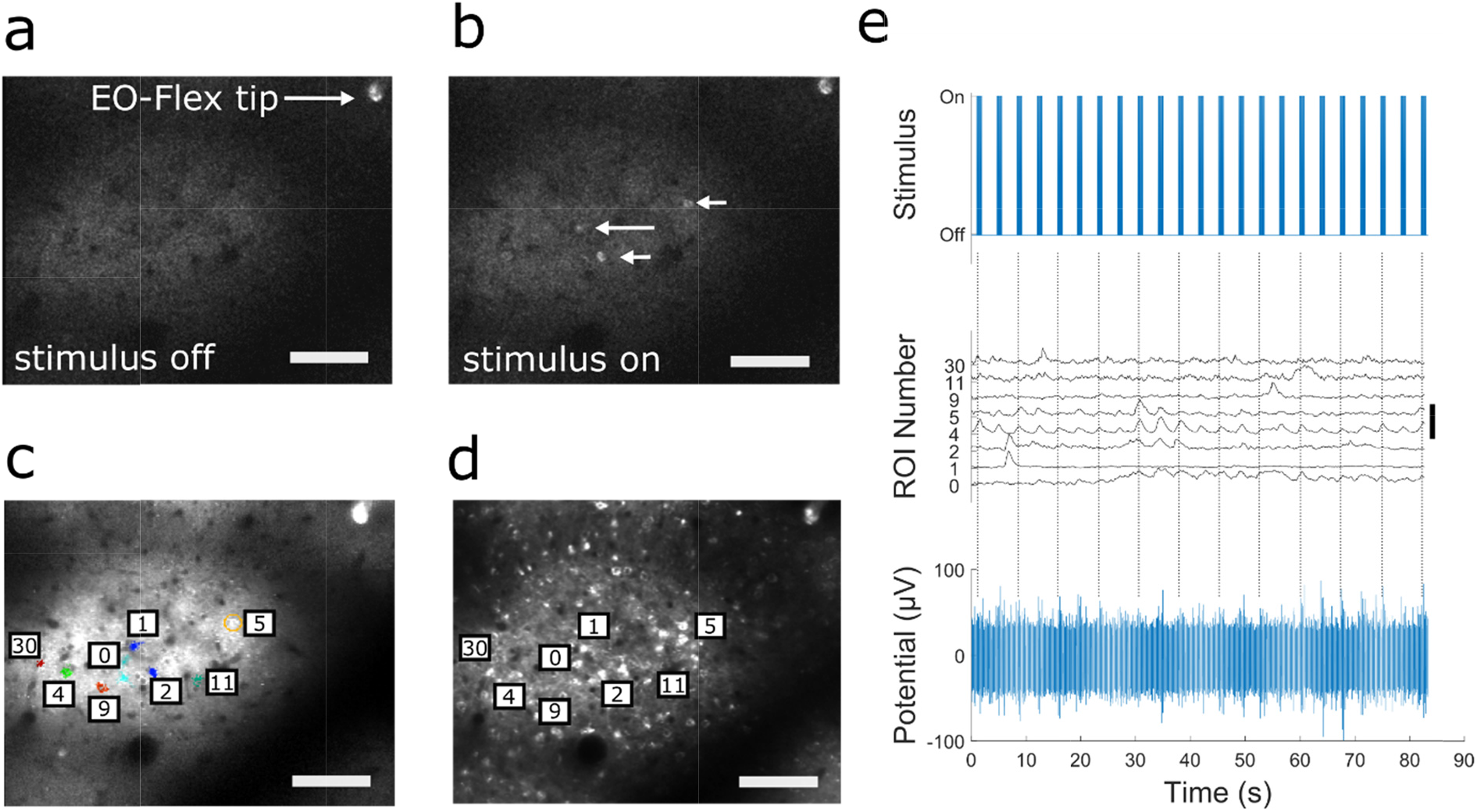
In vivo calcium imaging combined with simultaneous electrical recordings in the cortex of AAV2-CaMKII-C1V1-mCherry injected *Vglut2-GCaMP6f* mice confirms EO-Flex mediated optical excitation of neurons. (**a-b**) Fluorescence images from a time-lapse recording showing green fluorescent calcium indicator expressing neurons in layer 2/3 (depth, 270 μm) of *Vglut2-GCaMP6f* transgenic mice before (a) and immediately after (b) optical pulse train delivery to AAV2-CaMKII-C1V1-mCherry transduced cells. The probe tip is visible in the upper right corner of the field of view. The arrows in (b) indicate neurons that responded with fluorescence calcium transients to the optical pulses from the EO-Flex probe. Scale bar 60 μm. (**c**) Average fluorescence image showing GCaMP6f expressing cells overlaid with regions of interest (ROIs) active during the 80 s stimulation period. Active ROIs were identified by automated analysis of cellular calcium signals using Suite2p (see Methods). Scale bar 60 μm. (**d**) Average fluorescence image showing AAV2-CaMKII-C1V1-mCherry transduced neurons within the same field of view as in (c), confirming opsin expression in cells that show time-locked calcium responses to delivered optical pulses. Scale bar 60 μm. (**e**) Successful optical excitation of neural activity was confirmed by simultaneous electrical recordings with the EO-Flex probe. (top) Delivered optical pulses (stimulation frequency, 8 Hz; pulse width, 12.2 ms; on/off cycling, 1 Hz) using 600 μW of power at the probe tip. (center) Calcium transients within the individual ROIs indicated in panel (c). Scale bar represents 2 dF/F. (bottom) Recorded multi-unit activity after eliminating Becquerel effect mediated artifacts, caused by transient scanning of the imaging beam across the probe tip. Dashed lines are added for alternating pulse trains to aid in visual correlation between delivered optical pulses and measured calcium spiking and electrical activity.

**Supplemental Figure S9.**
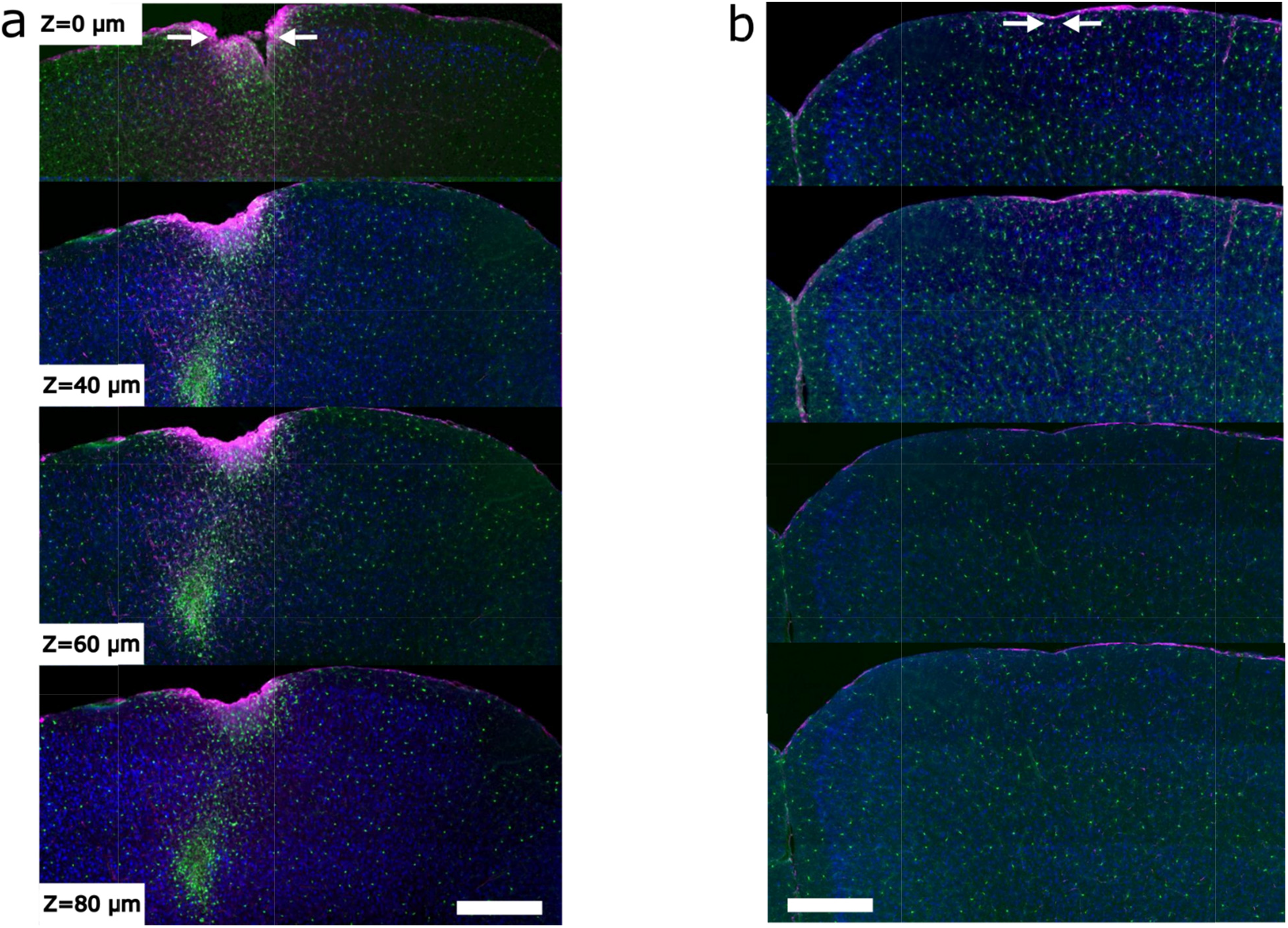
EO-Flex probes evoke minimal tissue inflammatory responses compared to multimode fibers commonly used in optogenetic experiments. (**a**)-(**b**) Example images of 20 μm thick serial coronal brain sections around the multimode fiber (a) and EO-Flex probe (b) implantation sites (boundaries indicated by white arrows). Both the multimode fiber (diameter, 250 μm) and EO-Flex probe (diameter, 12 μm) were advanced to ~1 mm depth into the cortex. Images were taken one week after brain implantation in *Cx3crl^GFP/+^* mice with labeled microglia (green). The sections were co-stained with anti-NeuN (blue) and anti-GFAP (magenta) antibodies to label neurons and astrocytes, respectively. z denotes slice spacing in microns. Scale bars 400 μm.

**Supplemental Figure S10.**
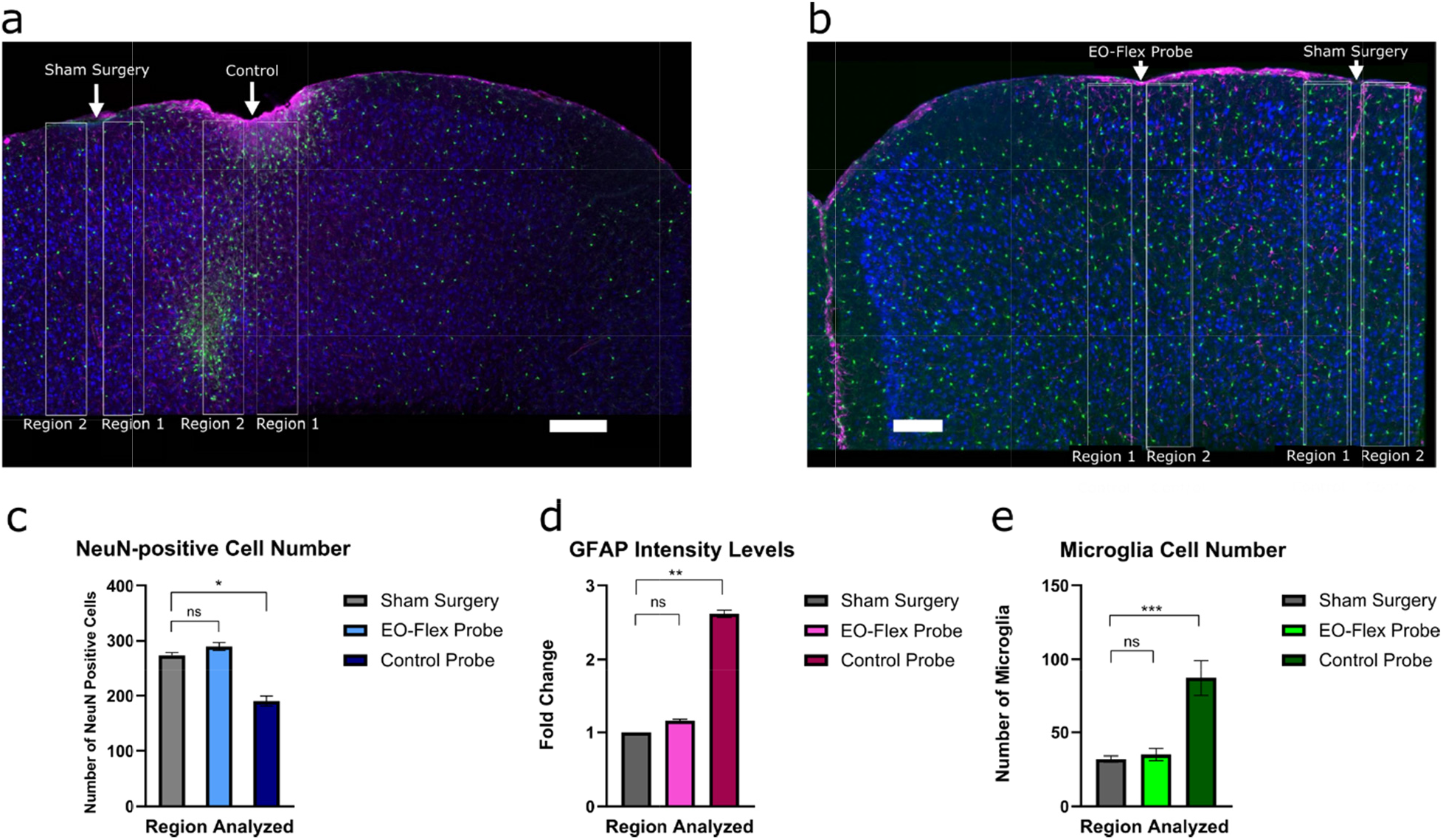
EO-Flex probes evoke minimal tissue responses compared to multimode fibers commonly used in optogenetic experiments. (**a-b**) Example images (from Fig. S9) showing the analysis approach. Cellular responses were quantified and averaged across two 150 μm x 1 mm analysis regions (white boxes) flanking each insertion site. To distinguish surgery from probe related tissue responses, an additional craniotomy of comparable size was made 0.7 mm lateral to each device implantation site. The same analysis approach was used to quantify cellular responses at this sham surgery site. Scale bar 200 μm and 150 μm respectively. (**c**) Population analysis showing the impact of multimode fiber or EO-flex probe implantation on neuronal cell numbers. (**d**) Population analysis showing the impact of multimode fiber or EO-flex probe implantation on astrocyte reactivity, as measured by GFAP expression level. (**e**) Population analysis showing the impact of multimode fiber or EO-flex probe implantation on microglia reactivity, as measured by microglial cell number.

**Supplemental Figure S11.**
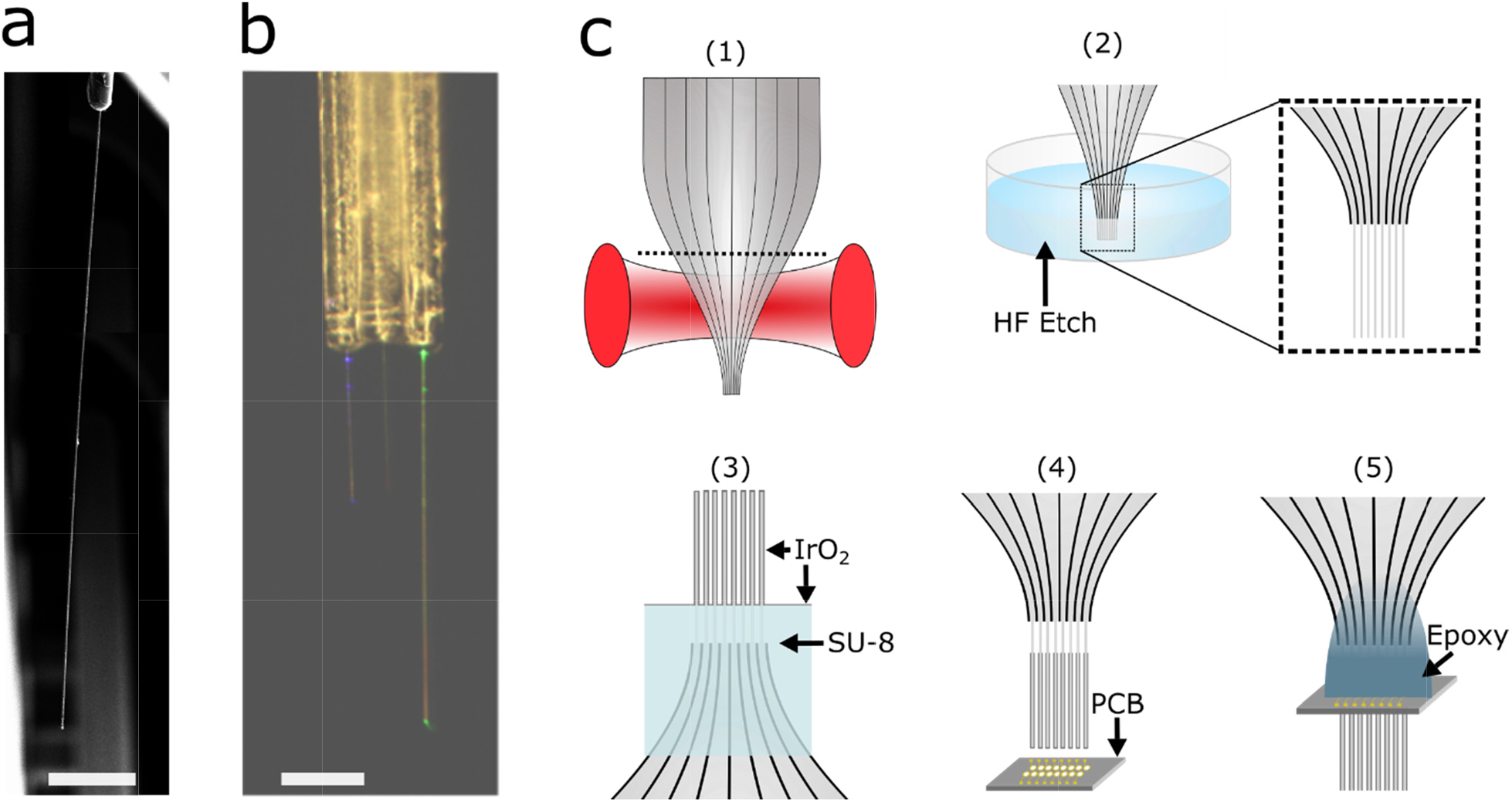
Scaling strategies of the EO-Flex probes for both probe length and arrays. (**a**) Two electron micrographs overlaid to show a 3.7 mm probe using a longer microfiber core. Scale bar 500 μm. (**b**) Image showing a 3×1 EO-Flex array with individually addressable optical channels (442nm and 543nm output). Scale bar 250 μm. (c) A proposed method for scaling probes into large two- or three-dimensional arrays (number of probes >100). (1) Utilize the same heat and pull strategy used to fabricate fiber bundles. Cut bundle at a desired backend diameter near the dashed line. (2) Use an acid bath to etch the surrounding cladding on the individual fiber cores with the length determined by the desired insertion depth. (3) Invert etched structure and mask the bottom portion of the device to ensure electrical channels remain separate, then deposit desired metal cladding layer(s). (4) Insert the array into a printed circuit board (PCB) with via spacings tuned for the desired probe spacing/density that also allows tolerance for the insertion process (e.g., ~50 μm). (5) After inserting the array into the PCB, individual electrical connections are made using for example pin-in-paste reflow soldering techniques. The final assembly would use an adhesive to form a stable mechanical interface between the fiber bundle and PCB. After which point the PEDOT:PSS and Parylene-C could then be deposited as previously described.

